# Adaptation to a Space-variant Visuomotor Delay can Cause Neglect-like Effects on Drawing Symmetry

**DOI:** 10.1101/193334

**Authors:** Chen Avraham, Guy Avraham, Ferdinando A. Mussa-Ivaldi, Ilana Nisky

## Abstract

In daily interactions, our sensorimotor system accounts for spatial and temporal discrepancies between the senses. Functional lateralization between hemispheres causes differences in attention and control of action. In addition, differences in transmission delays between modalities affects motor control. Studies on hemispatial neglect syndrome suggest a link between temporal processing and lateral spatial biases. To understand this link, we studied participants who performed lateral reaching, and adapted to delayed visual feedback in either left, right, or both workspaces. We tested transfer of adaptation to blind drawing, and found that adaptation to left or both delay caused selective leftward elongation. In contrast, adaptation to right delay caused elongation in both directions. Arm dynamics alone cannot explain these findings, but a model of a combined attentional-motor asymmetry across the hemispheres explains our observations. This suggests a possible connection between laterality in delay processing and motor performances observed in cases of hemispatial neglect.

## Introduction

When integrating external information for the execution of accurate hand movements, our sensorimotor system overcomes two challenges: laterality and time delays. Laterality is a result of the asymmetrical processing of workspace-specific information in the hemispheres (Reuter-Lorenz et al., 1990). Time delays are a result of sensory information transmission and processing time, and they may vary between modalities (Hopfield, 1995). Previous studies investigated how the sensorimotor system compensates for differences between the spatial representations in the workspaces (Heilman and Valenstein, 1979, Koch et al., 2011, Ziemann and Hallett, 2001), and for the delays between the different modalities (Miall et al., 1985, Miall and Jackson, 2006, Pressman et al., 2007, Nisky et al., 2008, Nisky et al., 2010, Nisky et al., 2011, Di Luca et al., 2011, Honda et al., 2012, Rohde et al., 2014). In this study, we use adaptation and transfer of adaptation paradigms to examine the interplay between these two compensatory processes.

Jordan and Rumelhart (Jordan and Rumelhart, 1992) suggested that the execution of accurate movements under various environmental conditions relies on the existence of forward models. A forward model is an internal representation of the environment that predicts the sensory consequences of a motor command; it is used in the control of movement, and it helps to compensate for changes in the sensory feedback during motor adaptation (Wolpert et al., 1995, Miall et al., 2007). In adaptation studies, the internal representation is typically evaluated from the movements of participants following visual or force perturbations. Throughout the adaptation, the participants modify the kinematics and dynamics of their movements to reduce task-related errors and to maximize task success (Krakauer et al., 2000, Cohn et al., 2000, Simani et al., 2007, Shadmehr and Mussa-Ivaldi, 1994). A common way to assess the adaptation and the construction of an internal model is by examining aftereffects when the perturbation is unexpectedly removed. Another approach is to test for transfer of adaptation to a different workspace (Shadmehr and Mussa-Ivaldi, 1994, Rotella et al., 2015), a different context (Kluzik et al., 2008), or a different task (Shadmehr and Mussa-Ivaldi, 1994, Botzer and Karniel, 2013). Investigating transfer of adaptation helps to understand how the new kinematics or dynamics are represented by the motor system.

One pathology that is associated with sensorimotor impairments in both laterality and temporal processing is *Hemi-spatial Neglect*, which occurs following stroke in one hemisphere, predominantly the right. Neglect patients demonstrate defective ability to report stimuli in the contralesional workspace (Colombo et al., 1976, Beis et al., 2004), and exhibit directional-motor impairments (Làdavas et al., 1993). Although most studies refer to neglect as spatial deficit, several studies also reported time-related impairments. For example, reports of a considerable delay in visual awareness of left stimuli compared to right stimuli (Robertson et al., 1998). This suggests that there is a link between laterality and temporal aspects of information processing.

In this study, we examined adaptation to a space-variant visuomotor delay– by considering the response to a 0.15 sec delay that affects selectively the visual feedback of hand movements in one direction (and consequently, one workspace). Spatially uniform visuomotor delay has been shown to cause alterations in movements’ extent (Botzer and Karniel, 2013). This study suggested that the sensorimotor system copes with delayed visual feedback by manipulating the current state variables, and specifically, by changing the gain in the internal representations. We hypothesized that by applying a delay exclusively in right-hand movements toward one workspace, and not toward the other, we can induce lateral asymmetry in the state representation of the arm, and find asymmetric aftereffects and asymmetric transfer of adaptation to different tasks. Taking into consideration only the known literature about visuomotor delay, we came up with two possible effects after exposure to workspace-dependent delay. It has been shown that adaptation to visuomotor gain has a wide range of generalization (Krakauer et al., 2000). Therefore, one possible prediction is that adaptation to workspace-specific delay will be generalized across space and cause elongation of movements toward both workspaces (Figure 1A, *Wide Generalization*). Second, the human brain sometimes learns context-depended perturbations (Howard et al., 2010). Therefore, if the brain was able to learn only the context of the workspace-specific delay, effects will be restricted to the delayed workspace (Figure 1A, *Narrow Generalization*).

**Figure 1.**
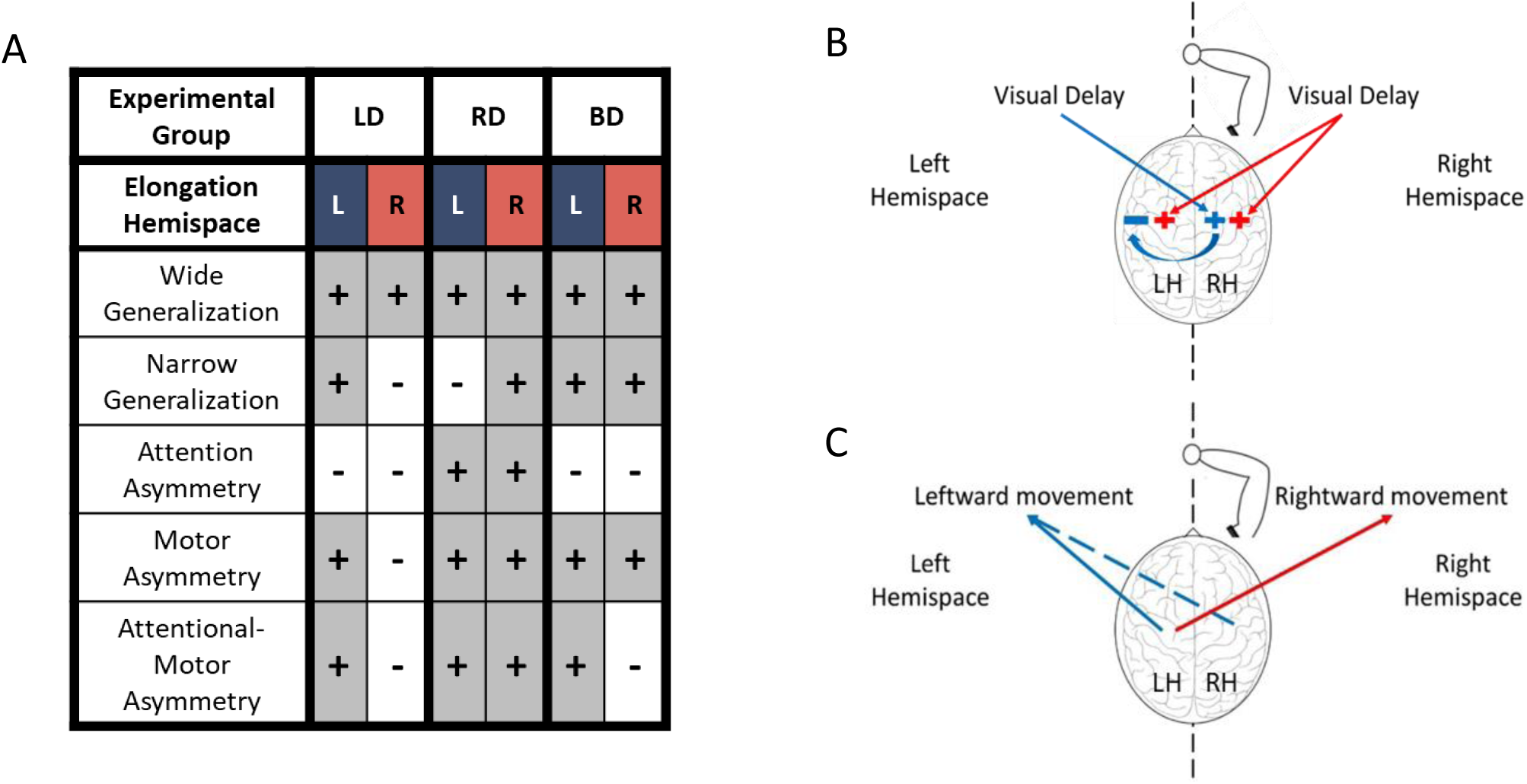
A model for the effect of delay in left, right, or both hemispaces on rightward and leftward movements. (A) Summary of all possible effects for hemispace-specific delay. For each condition of delayed hemispace, we expect movements to be elongated toward either the left or the right hemispaces, according to the accountable mechanism. (B) The effect of delayed visual feedback on the hemispheres according to the visual fields. Delay in left visual field (blue) affects motor circuits responsible for movement extension in the right hemisphere, and delay in the right visual field (red) affects both hemispheres. Following excitation of the right hemisphere after exposure to left delay, the right hemisphere inhibits motor circuits in the left hemisphere, thereby canceling any deviation toward the right hemispace after exposure to left delay (blue arrow). (B) The effect of the hemispheres on movement extent toward both hemispaces. The left hemisphere controls movements of the right hand toward left (blue) and right (red) sides, and the right hemisphere can mediate leftward movements (dashed blue).

However, it is possible for workspace-and direction-specific delay to have other effects on movements’ extent that would depend on the hemisphere that processes the visuospatial information. The hemispheres exhibit asymmetrical visuospatial attention. This is known as “right hemisphere dominance”, whereby the right hemisphere holds representations of both left and right fields (Heilman and Valenstein, 1979) and inhibits the left hemisphere (Ziemann and Hallett, 2001, Koch et al., 2011) and the left hemisphere processes only the right field (Figure 1B). Therefore, if the effect of delay on movement extent is solely attentional, when the delay is presented in the left workspace, it is predicted not to cause movements elongation in both workspaces, regardless of whether or not a delay is also presented in the right workspace; in contrast, for the right only delay case, movements will be elongated toward both workspaces (Figure 1A, *Attention Asymmetry*). Second, to account for laterality in the visuomotor control of the right hand in right-handers, studies suggested that the right hemisphere is involved in movements only toward the left workspace whereas the left hemisphere is involved in movements toward both right and left workspaces (Farne et al., 2003, Heilman and Valenstein, 2010). With this assumption of motor asymmetry (Figure 1C), left delay – visually processed in the right hemisphere – would affect movements’ extent only toward the left workspace, and right delay – visually processed by the left hemisphere – would affect movements’ extent toward both workspaces (Figure 1A, *Motor Asymmetry*). Lastly, if both hypotheses are correct, accounting for attention and motor asymmetry yields the following predictions: left only delay and right only delay would have the same effect as the *Motor Asymmetry hypothesis*; however, because of the attentional inhibitory circuit from the right to left hemisphere, presentation of both left and right delay will cause only leftward elongation (Figure 1A, *Attentional-Motor Asymmetry*).

To distinguish between the possible hypotheses, we asked participants to perform reaching movements to both left and right targets, and we applied a delay between the hand and the visual cursor in movements to one or both sides. We examined the effect of this delay on the amplitude of the reaching movements. To probe for laterality-related changes in the representation of the delay, we investigated the transfer of adaptation to a blind circle-drawing task, in which participants were requested to draw two-dimensional circles with multiple movement directions without visual feedback. We chose a blind drawing task because it allows for testing the effects of adaptation to delay when participants rely only on feedforward control and proprioceptive feedback, and because it allows for the detection of asymmetries in a continuum of directions (Punt et al., 2013). We found aftereffects of adaptation to delayed visual feedback in reaching movements, and transfer of adaptation to blind drawings. Interestingly, while the aftereffects reflected the spatial effect of the delayed perturbation, the transfer effects had significant asymmetries between delay conditions: only when the delay was presented in leftward reaches, regardless of whether it was also presented in the rightward reaches, participants exhibited asymmetrical neglect-like blind drawings. We account for these results with a computational model that includes an *Attentional-Motor Asymmetry* – laterality and right hemisphere dominance – together with an interplay between inverse and forward model adaptation with an unaltered endpoint stabilization controller.

## Results

### Experiment 1: Adaptation to a delay in one workspace

#### Reaching movements-Adaptation to delay affects reaching movements toward the delayed workspace

To examine the effect of delay that was presented only in one workspace and to distinguish between our hypotheses, two groups of participants took part in an adaptation experiment (Figure 2). They were asked to make center-leftward and center-rightward reaching movements with visual feedback of the cursor. The experiment consisted of Baseline, Adaptation, and Washout stages. During the adaptation stage and only in the leftward (left delay, LD, group, N=15) or rightward (right delay, RD, group, N=15) reaching, the continuous visual feedback of the cursor was delayed by 0.15 sec. The visual feedback was presented without delay for movements to the other direction and in the other stages of the experiment. To assess the effects of delay and adaptation, we examined movement extent in reaching.

**Figure 2.**
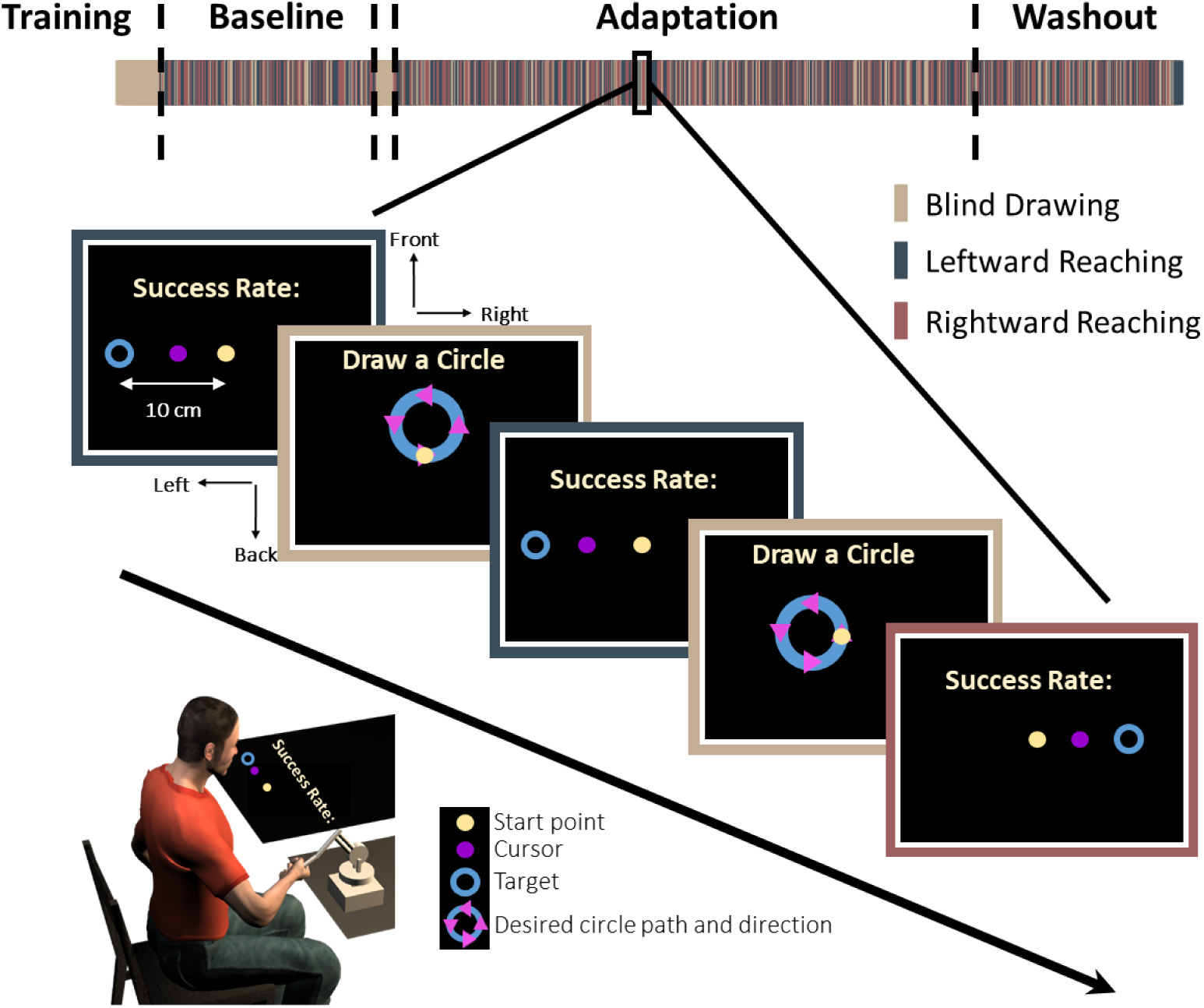
Experimental protocol. In each trial, participants were required to make a *reaching*: move a cursor between a start and an end target to the left (blue bar) or the right (red bar), or to make a *blind drawing*: draw a circle without visual feedback (beige bar). In the reaching, a start point (light yellow), a target (blue circle), and a cursor were presented. In the blind drawing, a desired circle path (blue) was presented together with arrows that indicated movement direction (magenta triangles), but no cursor was presented. Overall, there were eight different kinds of circular movements: four different locations – front, back, right and left, and two different directions – clockwise and counterclockwise. The experiment was divided into three sessions: Baseline, Adaptation, and Washout. During the Baseline and Washout sessions, the cursor movement in the reaching task was concurrent with the movement of the hand. During the Adaptation session, the visual feedback was delayed by 150ms in movements towards the leftward, the rightward, or both targets (see Methods for details).

Reaching movement analyses of the left and right delay group suggest that both groups adapted to the delay (Figure 3). Upon early exposure to the delay (EA), participants over-reached the target in the workspace to which the delay was applied. With repeated exposure to the perturbation, they quickly adjusted their movements, and by the late stage of adaptation (LA), they restored Late Baseline (LB) performance. They also initially started to under-reach the target in the opposite direction, but this effect was weaker and vanished quickly. After the delay was removed (EW), we observed an aftereffect of target under-reach only in movements toward the delayed workspace.

**Figure 3.**
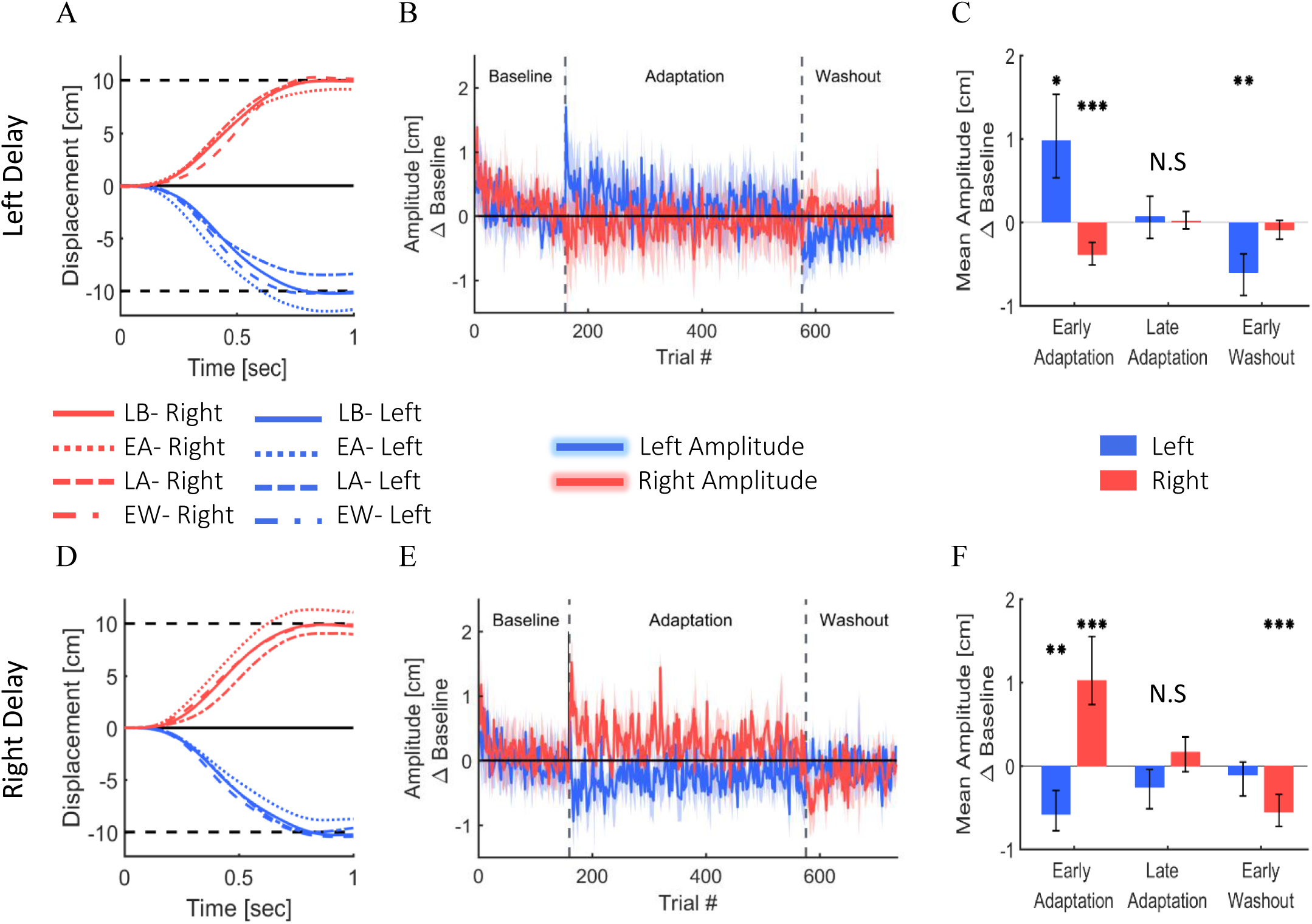
Reaching movements from the left delay (A-C) and right delay (D-F) conditions of Experiment 1. (A) Examples of movements of a typical participant in the Left Delay group from the Late Baseline, Early adaptation, Late adaptation, and Early Washout stages. Positive displacement indicates a rightward movement. The participants overshoot the left target when initially exposed to delay, but they quickly adapt and restore baseline movements, and exhibit undershoot in the washout. Interestingly, the movements in the other direction are initially affected, but no aftereffects are observed. (B) Amplitude (line) and 95% confidence intervals (shaded region) of the leftward and rightward movements from the Left Delay condition. Results are presented after subtraction of the movement amplitude at the end of the baseline session and taking absolute value. Positive (negative) value indicates overshoot (undershoot) in the direction of movement. Leftward movements demonstrate typical pattern of adaptation, and the rightward movements exhibit an initial undershoot that is reduced with adaptation and no aftereffect. (C) Mean Amplitude in the presence of left delay in the first and last five movements of the Adaptation stage and the first five movements of the washout for all participants. (D-F) Similar but mirror results were observed in the Right Delay condition.

These observations were supported by a statistical analysis. Within each experimental group, we found significant changes in the movement amplitude between the different stages in the experiment, and these changes were different between left and right movements (Stage-Workspace interaction effects – LD: F_0.87,12.15_=95.14, p<0.001; RD: F_3,42_=45.92, p<0.001). In the leftward reaches of the left delay group, we observed a typical adaptation pattern: overshoot in EA (t_14_=3.59, p<0.05); no difference in LA (t_14_=0.48, p=1); and undershoot in EW (t_14_=4.53, p<0.01) (all with respect to LB, Figure 3C). The rightward reaches of this group exhibited a different pattern: undershoot in EA (t_14_=5.92, p<0.001); and no difference in LA (t_14_=0.27, p=1) and EW (t_14_=1.53, p=0.88). A similar but opposite pattern was observed in the right delay group (rightward reaches: EA: t_14_=5.22, p<0.001; LA: t_14_=1.57, p=0.83; EW: t_14_=5.47, p<0.001; leftward reaches: EA: t_14_=4.83, p<0.001, LA: t_14_=2.01, p=0.38, and EW: t_14_=1.14, p=1, Figure 3F).

Overall, in both the left and right delay groups, the participants adapted to the workspace-specific visuomotor delay by selectively adjusting their movement amplitude, and exhibited significant aftereffects of adaptation in the workspace where the delay was applied. The initial undershoot to the other workspace during EA quickly vanished, and there were no aftereffects in the non-delayed workspace.

#### Blind drawing task-Transfer of adaptation causes spatial asymmetry that depends on the delayed workspace

To test the transfer of adaptation, we examined the symmetry of blind circle drawing movements that were interleaved with reaching movements. Participants drew left, right, front, and back circles in clockwise (CW) and counterclockwise (CCW) directions without any visual feedback. To assess the symmetry, we calculated the left and right error relatively to an ideal circle.

In both LD and RD groups, the transfer of adaptation yielded a clear spatial elongation in the blindly drawn circles. However, in a striking contrast to the effects of left and right delay on the reaching movements, the patterns of elongation differed substantially between the two groups. An example of drawings following adaptation to left delay is depicted in Figure 4A. Following adaptation to left delay, the participants drew left-initiated circles that were elongated to the left, whereas the right-initiated circles were not elongated at all. This was also evident in the group analysis (bar plots Figure 4A and Figure 4C). In contrast, following adaptation to right delay, participants drew both left- and right-initiated circles that were elongated to the direction of their initiation – left-initiated circles were elongated to the left, and right-initiated circles were elongated to the right (Figure 4B,E). The effect of the initiation workspace is especially highlighted in the front and back circles: the side of the elongation is determined by the CW and CCW drawing direction (orange and green traces, respectively) rather than by the spatial location of the circle. The effect of delay on the circle drawing task persisted also to the washout stage. This was despite the fact the extents of the reaching movements returned very quickly to those observed in the Baseline.

**Figure 4.**
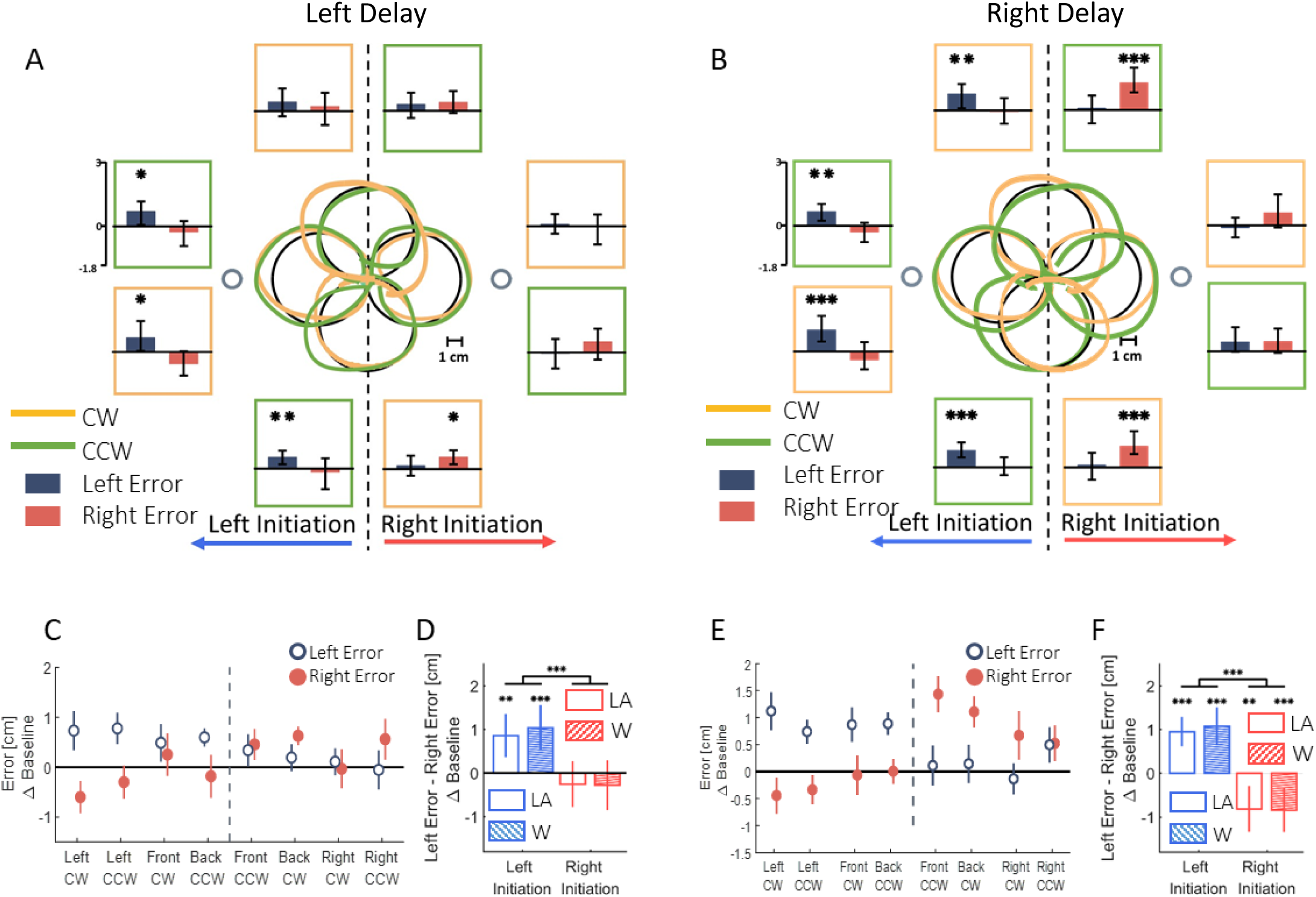
Left and right deviation from the desired trajectory in the circle drawing task in Experiment 1. (A) At the center, examples of individual movements of a typical subject that illustrate the deviation of the drawn circles, for both clockwise (orange) and counterclockwise (green) circles. Large black circles are the ideal drawings, and the two small circles are the targets from the reaching task (drawn at scale). Panels around the center present mean difference of left (dark blue) and right (dark red) error for circles drawn in the end of adaptation session in the presence of delay only in the left side of the tasks space. The panels are located spatially to represent the location and drawing direction of the circles. (B) Similar to A following adaptation to a delay only in the right side of the tasks space. (C) Left and right error following adaptation to Left Delay as a function of the location (front, back, right and left) and the direction (clockwise – CW – and counterclockwise – CCW) of the drawn circle. The dashed line divides the circles to left- and right-initiated circles. The elongation is observed only in the left side of the left-initiated circles. (D) Statistical analysis of the difference in left and right error in the Left Delay group shows deviation only toward the left side. Empty bars are for Late Adaptation session and bars with stripes are for Washout session. (E) and (F) are similar to (C) and (D) but following adaptation to Right Delay. Surprisingly, the result of the Right Delay is not a mirror picture of the Left Delay condition. Instead, both left- and right-initiated circles are elongated in the side of their initiation hemispace. All error bars are bootstrap 95% confidence intervals.

To highlight the laterality of the spatial effects, we performed a summarizing analysis, in which we distinguish between the circles based on the workspace of the initial drawing movement: left-initiated circles are left, front CW, and back CCW, and right-initiated circles are right, front CCW, and back CW. Then we calculated the difference between the left and right errors within each group (Figure 4D and 4F). In the LD group, we found a significant change in the elongation of the circles between the stages (Workspace-Stage interaction effect: F_0.97,57.23_=18.14, p<0.001). Specifically, the left errors were significantly larger than the right errors (meaning left elongation) only for the left-initiated circles during both Late Adaptation (LA: t_59_= 3.47, p<0.01) and Washout (W: t_59_= 3.96, p<0.001). In the RD group, we also found a significant change in the elongation of the circles between the stages (Workspace-Stage interaction effect: F_0.96,57_=76.44, p<0.001). However, following adaptation to right delay, the left errors were significantly larger than the right errors in left-initiated circles (LA: t_59_=6.74, p<0.001; W: t_59_=4.83, p<0.001) and right errors were significantly larger than the left errors (meaning right-elongation) in right-initiated circles (LA: t_59_=3.29, p<0.01; W: t_59_=4.17, p<0.001, see Figure 4F).

From Experiment 1 we conclude that after adapting to a visuomotor delay between the movement of the hand and its visual feedback in either the left or the right workspaces, participants present aftereffects in reach movements to the workspace in which the delay was presented, consistent with context-depended adaptation. They also exhibit transfer to blind drawing that causes spatial elongation of the drawing, and the pattern of elongation along the frontal plane depended on the workspace in which delay was presented – left delay caused asymmetrical elongation only to left initiated circles and right delay caused symmetrical elongation to both left and right initiated circles. This is in agreement with the expected results for both *Motor Asymmetry* and *Attentional-Motor Asymmetry* hypotheses. This shows that exposure to delay might be processed differently according to the workspace in which it was presented, and that the laterality in the visual feedback is important for shaping our representation of the environment when adapting to temporal misalignment between the different sensory streams.

#### Experiment 2: Adaptation to a delay in both workspaces

To distinguish between *Motor Asymmetry* and *Attentional-Motor Asymmetry*, we performed another experiment in which the delay was presented in both workspaces (both delay, BD, group, N=20). In addition, a control group performed the experiment without any perturbation (no delay, ND, group, N=15).

#### Reaching task-Adaptation affects movements toward both workspaces

The extent of reaching movements for the both delay group demonstrated a typical pattern of adaptation that was similar in both directions (Figure 5A-C). There was a statistically significant difference in movement extent between the stages (Stage-F_1.55,29.26_=60.51, p<0.001), but no difference between leftward and rightward movements in the different stages (Direction-F_0.51,9.75_=2.78, p=0.13 and Direction-Stage interaction-F_1.55,29.26_=2.38, p=0.12). When the delay was first introduced, movements over-reached the target in both sides (t_19_=4.27, p<0.01). Continued exposure to delay in both workspaces led to reduction of the over-reaching pattern, though the adaptation was not fully achieved compared to baseline performances (t_19_=3.11, p<0.05). When the delay was removed, participants under-reached the target in both sides (t_19_=7.74, p<0.001). These results indicate that when the visual feedback is delayed in both workspaces, the participants adapt to the perturbed visual feedback, and exhibit aftereffects in both workspaces.

**Figure 5.**
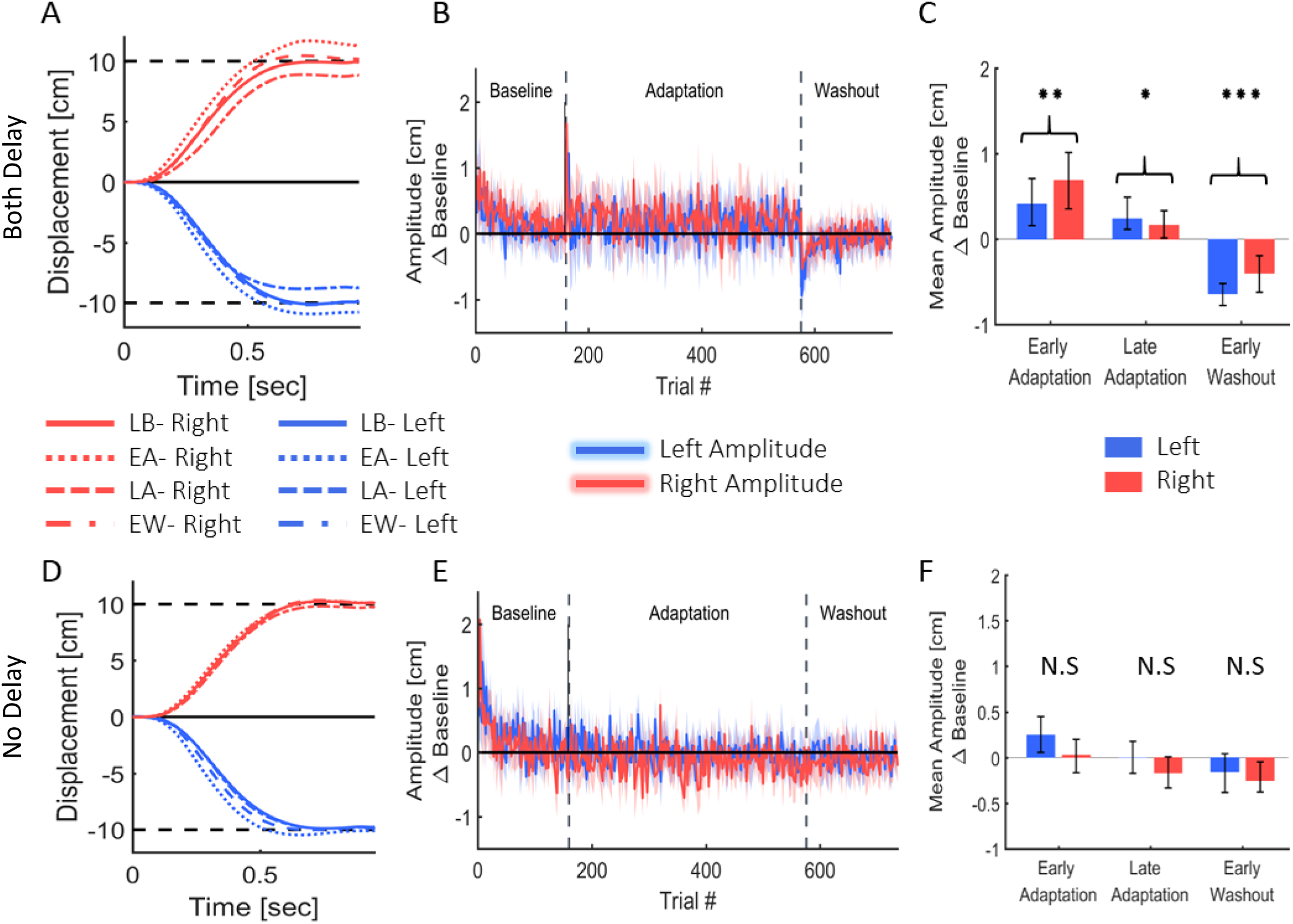
Reaching movements from the Both delay (A-C) and No delay (D-F) conditions of Experiment 2. (A) Examples of movements of a typical participant in the Both Delay group from the Late Baseline, Early adaptation, Late adaptation, and Early Washout stages. Positive displacement indicates a rightward movement. The participants overshoot both targets when initially exposed to delay, but they quickly adapt and restore baseline movements, and exhibit undershoot in the washout. (B) Amplitude (line) and 95% confidence intervals (shaded region) of the leftward and rightward movements from the Both Delay condition. Results are presented after subtraction of the movement amplitude at the end of the baseline session and taking absolute value. Positive (negative) value indicates overshoot (undershoot) in the direction of movement. (C) Mean Amplitude in the presence of delay in both sides in the first and last five movements of the Adaptation stage and the first five movements of the washout for all participants. (D-F) Results for the No Delay condition. Graphs and colors are as in (A-C). No spatial deviation is observed, as expected.

The control group did not experience any visual perturbation (Figures 5D-F), and did not demonstrate any deviation in movement extent. This corroborates our claim that the observed spatial deviations are a result of the delayed visual feedback.

#### Blind drawing task-transfer of adaptation affects only the left side of the drawn circles

The results of the transfer of adaptation to the blind drawing task in the BD group were very similar to those of the left delay group, and hence, consistent with the *Attentional-Motor Asymmetry* hypothesis. We found a significant main effect of initiation workspace, stage, and the interaction between stage and initiation workspace (F_0.54,42.75_=226.45, p<0.001, F_1.1,85.5_=7.8, p<0.01, and F_1.1,85.5_=11.68, p<0.001, respectively). Even though the delay perturbation was presented in both sides, only the left-initiated circles were elongated to the left (Figure 6A-B, positive difference between left and right error compared to the baseline difference LA: t_79_=4.75, p<0.001; W: t_79_=3.65, p<0.01), and the right-initiated circles were not elongated at all (LA: t_79_=0.23, p=1; W: t_79_=0.25, p=1).

**Figure 6.**
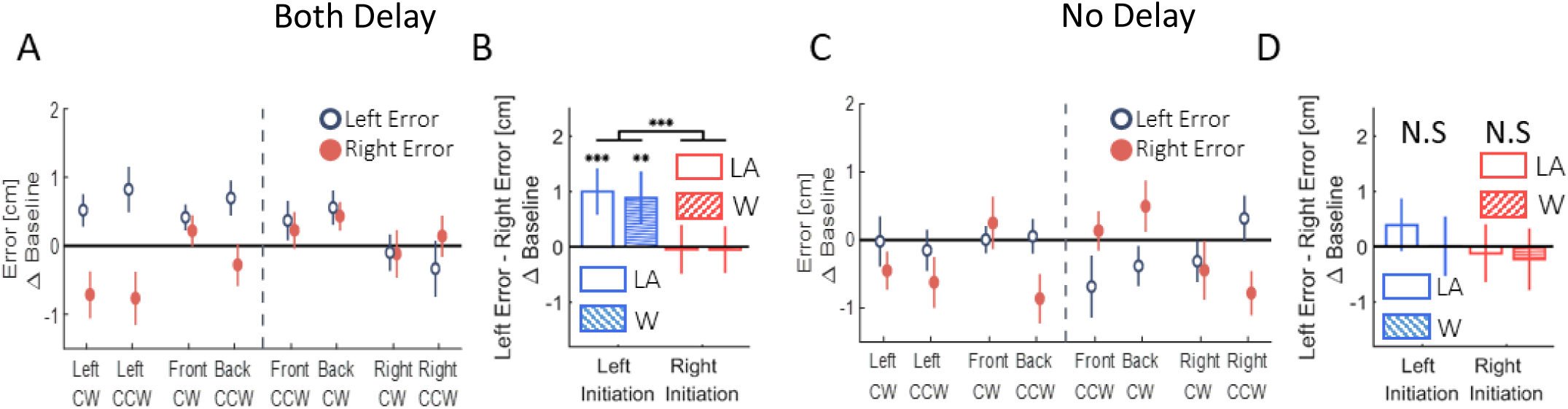
Left and right deviation from the desired trajectory in the circle drawing task in Experiment 2. (A) Both delay condition. Left and right error following adaptation to delay in both hemispaces as a function of the location (front, back, right and left) and the direction (clockwise – CW – and counterclockwise – CCW) of the drawn circle. The dashed line divides the circles to left- and right-initiated circles. The error is different according to the side where the drawing is initiated: when the drawing is initiated in the left– left error is larger than right error, and when the circles are initiated in the right– no deviation is observed. (B) Statistical analysis of the difference in left and right error in Both delay condition. Empty bars are for Late Adaptation session and bars with stripes are for Washout session. The graph show a spatial deviation to the left side, when the circles are initiated in the left. (C) and (D) are similar to (A) and (B) but for No delay condition. No similar pattern of difference between left and right error is observed.

In the control experiment, with no perturbation (Figures 6C-D), the circles were nearly symmetrical without any lateral pattern. This corroborates that the elongation of the blind circles is not caused by unrelated effects of our setup or fatigue. We performed another control analysis on the drawings of participants from all four conditions (LD, RD, BD, and ND) – we calculated the front and back deviation from ideal circles. There were no consistent elongation to the front and to the back of neither right- or left-initiated circles (Figure 1 in supplemental information), suggesting that the transfer effect was specific to the lateral dimension of movement.

The laterality in the effect of adaptation to delay provides an important insight on the laterality in the brain and the mutual influence between the two hemispheres. In contrast to the right delay group of Experiment 1, here, when participants adapted to delayed visual feedback in the right workspace concurrently with delay in the left workspace, their right-initiated drawings were not affected. These can indicate that activation of right hemisphere because of exposure to delay in the left workspace could have an inhibitory effect on the activation in the left hemisphere. Accordingly, the observed results are an evidence for *Attentional-Motor Asymmetry* in the hemispheres.

#### Simulation study – a model that includes asymmetrical transfer across hemispheres is critical

Taken together, the results of both experiments indicate asymmetry in the transfer of adaptation. These asymmetries indicate that the processing of neural information is different between the hemispheres. However, there is also the alternative that simple effects of biomechanics could explain them. To address this question, we simulated the effects of exposure to delay, adaptation, and its transfer to blind circle drawing using a simplified computational model that is depicted in Figure 7. The arm was simulated as a two link planar kinematic chain (Pressman et al., 2008, Nisky et al., 2011). The controller included a trajectory and an end-point controller (Scheidt and Ghez, 2007, Botzer and Karniel, 2013). The trajectory controller consisted of a feedforward controller – an inverse model of the arm, and two feedback controllers - for vision and for proprioception (Ghez et al., 2007, Scheidt and Ghez, 2007, Scheidt and Stoeckmann, 2007, Scheidt et al., 2011), and received as an input a desired trajectory. The endpoint controller was implemented as a spring and a damper with an equilibrium at the desired static end of movement, and it stabilized the arm at the end of movement. The effect of adaptation in both reaching and circle-drawing was on movement’s extent, and therefore, and for simplification, we used a gain (G) to simulate the representation of delay in the feedforward controller, and the corresponding inverse of the same gain (1/G) in the visual feedback controller. We assumed that the proprioception does not change during the experiment, and therefore, the gain of the proprioceptive feedback was never modified.

**Figure 7.**
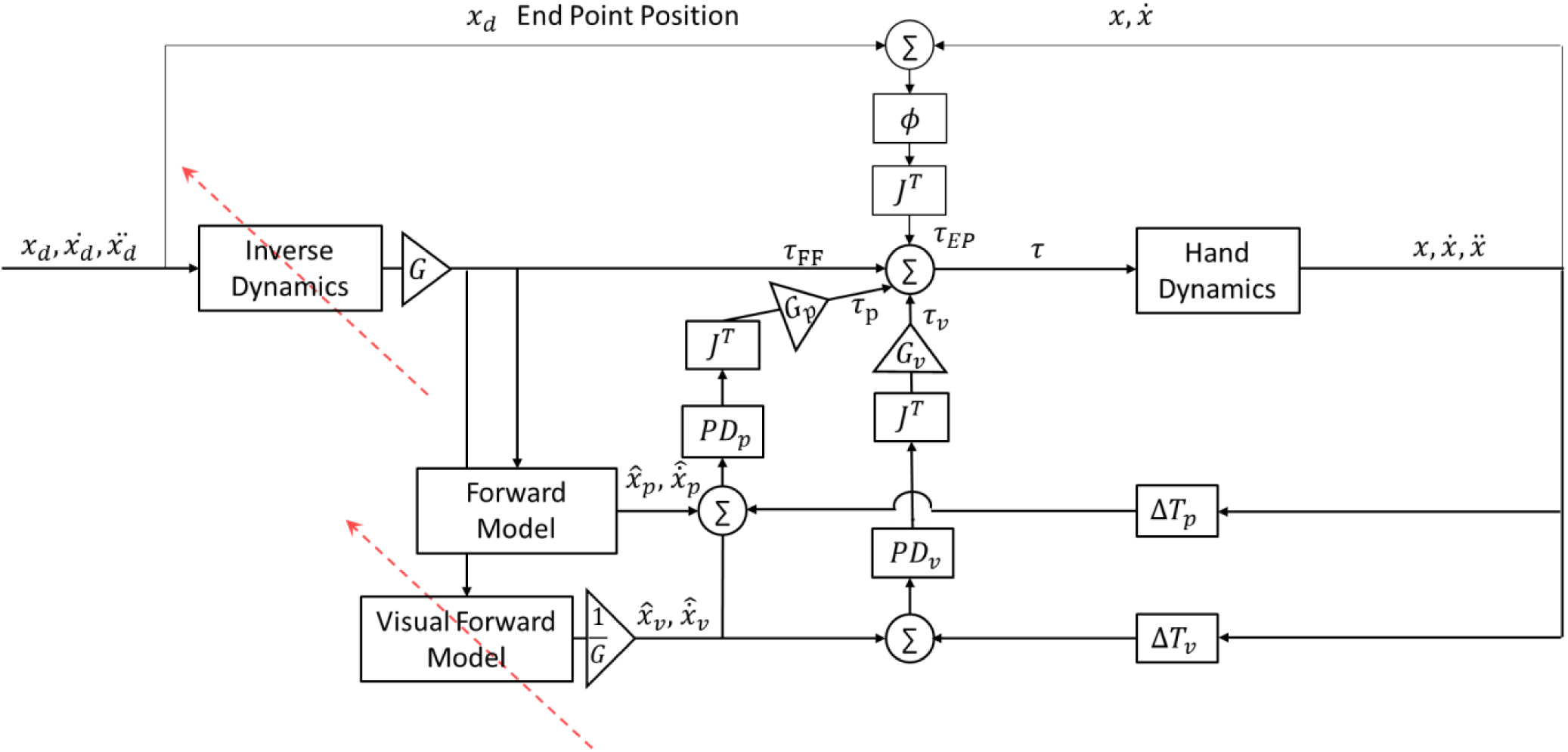
Simulation of hand movement with feedback and feedforward controllers. The desired torques are computed using the error between the estimated and the actual location and velocity of the hand. Additional end-point controller is used in order to reduce the error between the actual location of the hand to the desired end point.

Using this simulation of arm dynamics, we simulated the results that were observed in the different stages of the experiment. In the Baseline session, no perturbation was applied, and the simulated arm followed the desired trajectory properly (Figure 8A, solid line). During the Early Adaptation session, the visual feedback was delayed, but no change in the gains of the feedforward or feedback controllers has occurred yet. Hence, a misalignment between the estimated location and the actual observed location of the hand during the reaching task resulted in a positive error, and the feedback controller of the visual modality caused target over-reaching (Figure 8A, dotted line). In contrast, the vision-omitted movements were not hypermetric.

**Figure 8.**
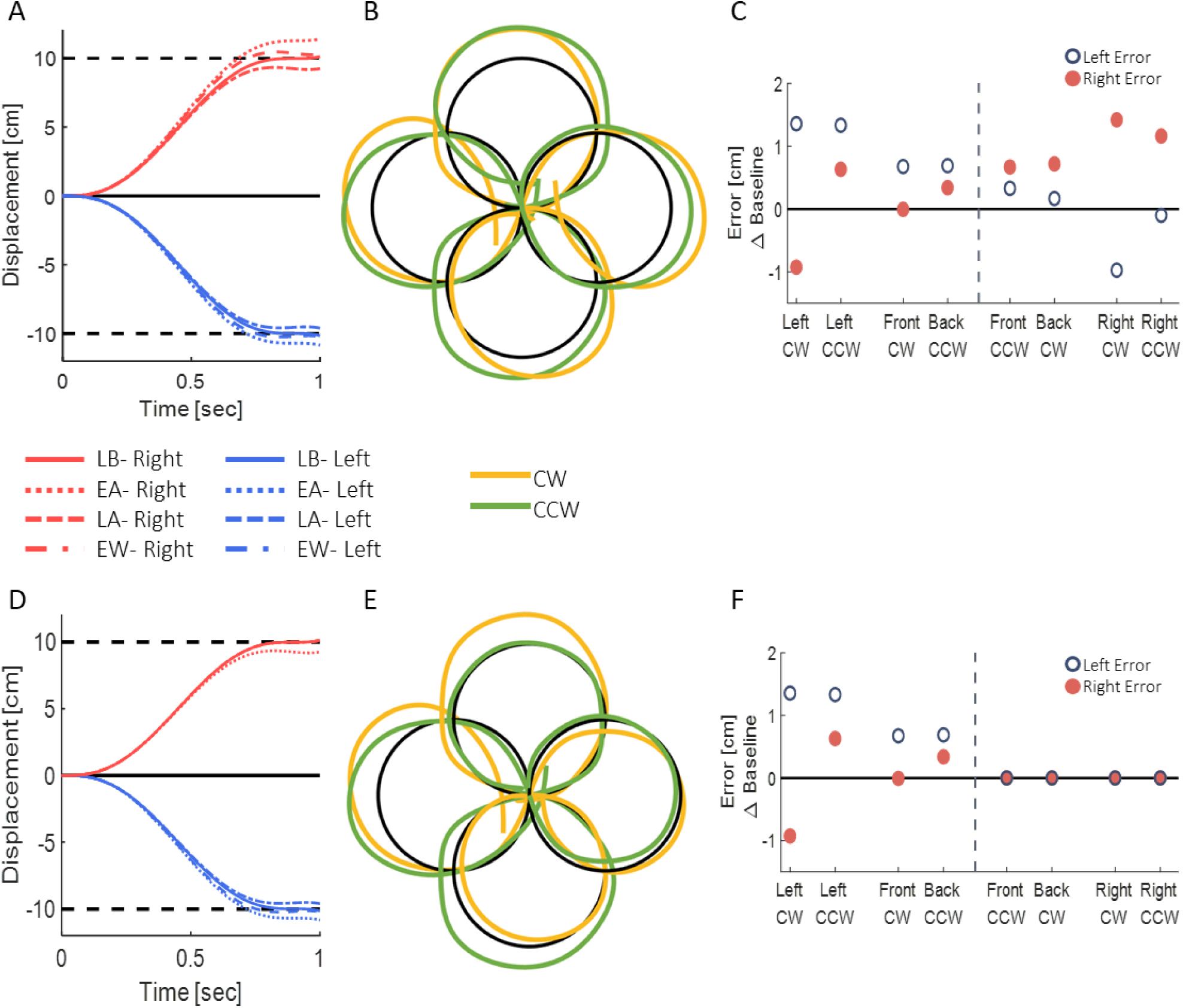
Simulation results without (A-C) and with (D-F) context-depended adaptation and transfer of adaptation. (A) Simulated reaching movements in the different stages of Late Baseline, Early adaptation, Late adaptation, and Early Washout. Positive displacement (red) indicates a rightward movement. Movements in both directions demonstrate typical pattern of adaptation to delay. (B) Simulated blind circles after adaptation to delay for circles in clockwise (orange) and counterclockwise (green) directions. The circles are elongated toward the hemispace where the drawing was initiated. (C) Left and right error as a function of the location (front, back, right and left) and the direction (clockwise – CW – and counterclockwise – CCW) of the drawn circle. The dashed line divides the circles to left- and right-initiated circles. Both left- and right-initiated circles are elongated in the side of their initiation hemispace. Using arm dynamics solely, we could not simulate the asymmetrical elongation pattern observed in our results. (D) Simulation of reaching movements in the presence of delay only in the left hemispace with context-depended adaptation. Movements toward the left target demonstrate typical pattern of adaptation, and movements toward the right target have only initial aftereffect when first exposed to the delay. (E) and (F) are similar to (B) and (C) but with context-depended transfer of adaptation. Adding the context-depended learning, we simulated the asymmetrical pattern of adaptation observed in the Left and Both delay groups.

After adapting to the delay, the movements with visual feedback gradually returned to baseline. To simulate this adaptation to delay, we used gain as representation of delay. We simulated the post-adaptation condition as magnifying gain (G>1), multiplied by the output of the forward model. Meaning, that the desired trajectory was extended in the direction of the movement. In the reaching movement with visual feedback, the visual forward model was multiplied by the inverse gain, causing a reduction of the error in the visual feedback controller, and leads to reduction of the over-reaching pattern (Figure 8A, dashed line). In the circular movements, the feedforward controller was multiplied by magnifying gain and cause elongation of the trajectory in the direction of movement. However, the end-point controller is activated at roughly the middle of the movement and returns the hand to the desired end-point. Therefore, the elongation is only observed in one side of the circle as depicted in Figure 8B,C. Throughout the movement, proprioception was not affected, and the feedback controller of this modality was only tuned to move the hand in the desired trajectory.

Following abrupt removal of the delay, the forward and inverse models were still tuned to the delayed condition. However, the visual feedback matched the real location of the hand, which resulted in negative error of the visual modality and under reaching of the target (Figure 8A, dashed and dotted line). This changes in the visual feedback did not affected the circles in the post adaptation condition (Figure 8B,C).

From this simulation, we saw that arm dynamics and representation of delay as visuomotor gain can explain the adaptation pattern in the reaching task, and the elongation of the blind drawn circles. However, the adaptation in the reaching task was in both leftward in rightward movements, similar to the adaptation in the Both Delay group (Figures 5A and 8A). Additionally, the elongation in the blind circles was toward both initiated workspaces, similar to the drawn circles observed in the Right Delay group (Figures 4B,E and 8B,C). Meaning, the asymmetry observed in our experiment could not be simulated with a uniform gain across directions. Therefore, a context-depended mechanism is necessary to simulate the workspace-specific effects. By adding an *Attentional-Motor Asymmetry* adaptation, we were able to simulate the reaching movements observed for the groups that were exposed to delay only in one workspace (Figures 3A and 8D), and the circles observed in the Left and Both Delay groups (Figures 4A,C and 8E,F).

## Discussion

In this study, we set out to establish the link between spatial representation of information across workspaces and adaptation to temporal misalignment between the senses. We investigated the effect of delayed visual feedback of cursor movement that is presented exclusively in one or in both workspaces on the movements of participants with and without visual feedback. We show that following an exposure to a visuomotor delay either in one or both workspaces, participants modified the extent of their reaching movements: the abrupt presentation of the delay caused hypermetria – participants made larger reaching movements; they reduced this hypermetria throughout adaptation, and exhibited aftereffects in the workspace where the delay was applied. This means that to reduce the overshoot of the target, participants compensated for the changes in the visual feedback by constructing an internal representation of the perturbation that was specific to the workspace it was applied in. Importantly, the effects of workspace-specific delay in the left and right workspaces mirrored each other.

In contrast, transfer of adaptation to the blind circle-drawing task revealed a different picture. Following adaptation to visuomotor delay, we observed hypermetric circles that were elongated only in one side. Whether the circles were hypermetric depended on the workspace of initiation of the drawing (left or right) and on the workspace in which delay was presented. The effect of the workspace of drawing initiation on the side of the circle that was hypermetric was demonstrated most clearly in the circles that were drawn in the front and the back locations. Although these circles were all in the middle of the task space, the drawings were different depending on the workspace where they were initiated. Interestingly, the hypermetria in the drawings was different between the left delay, right delay, and both delay groups. Adaptation to left delay or delay in both workspaces caused elongation of only leftward blind drawings. In contrast, adaptation to right delay caused elongation in both directions. A simulation study confirmed that arm dynamics alone cannot explain these findings. Instead, we had to include an asymmetrical, workspace-dependent, transfer of adaptation. We concluded that visuomotor delay might be processed differently depending on the workspace in which it was presented, and we further suggest that this difference resulted from *Attentional-Motor Asymmetry* between the hemispheres.

### Right hemisphere dominance and a model for laterality in the processing of visuomotor delay

When faced with an imbalanced stimulation across space, the hemispheres demonstrate different patterns of activation and inhibition, and these are reflected in asymmetric attention, perception, and action across workspaces (Reuter-Lorenz et al., 1990). An example of an asymmetric attention in healthy individuals is leftward perceptual bias – a spatial deviation toward stimuli located on the left side. This bias was suggested to arise from asymmetries in hemispheric activation: left hemisphere is activated only by stimuli in the right hemispatial field, while the right hemisphere is activated in response to stimuli in both the left and the right hemispatial fields (Heilman and Valenstein, 1979). In addition the right hemisphere can also interact more strongly with the left hemisphere, by exerting inhibition activity over cortical areas in the left hemisphere (Koch et al., 2011, Gotts et al., 2013). Because the activation process in the right hemisphere occurs in different locations for right or left stimuli (Corbetta et al., 1993), it is possible that the inhibition activity from right to left will only take place in response to left stimuli. Regarding to the control of right hand movements in right handers, it is well established that the left hemisphere controls movements toward both workspaces. However, the right hemisphere can contribute to the control of movements toward the contralateral space with the right hand (Farne et al., 2003, Heilman and Valenstein, 2010). This explains why in the case of processing delayed visual feedback in our experiment, leftward movements with the right hand can be strongly affected also by the right hemisphere.

Based on these patterns of activation and inhibition, we suggest that exposure to delay excites motor circuits associated with movement extension in the relevant hemisphere, such that: (1) Delay only in the left workspace has excitatory effect on brain areas responsible for movement extension in the right hemisphere (Figure 1B). Therefore, exposure to delay only in the left visual field causes only leftward hypermetria (Figure 1C). (2) Delay in the right workspace affects both hemispheres (Figure 1B), resulting in transfer of hypermetria toward both workspaces (Figure 1C). (3) Delay in both workspaces excites motor areas in both hemispheres. However, as a result of exposure to left delay, the right hemisphere inhibits the left, and cancels the excitatory effect of delay (Figure 1B). Overall, excitation effect is only maintained in the right hemisphere, thereby affecting leftward movements performed without visual feedback and causing leftward hypermetria.

### Adaptation and representation of visuomotor delay

Visuomotor delay was investigated in various types of movements, such as driving (Cunningham et al., 2001), tracking (Foulkes and Miall, 2000, Leib et al., 2017), and reaching (Botzer and Karniel, 2013). However, the effect of workspace-specific visuomotor delay was not investigated. One exception is a recent study in which participants were exposed to visuomotor delay while performing a complex task of Pong game in one side of the task space. The effect of the delay was examined by reaching movements with no visual feedback performed at the other side. The results of this study showed asymmetrical generalization from left to right but not from right to left (Farshchiansadegh, 2012). We also found evidence for initial generalization in the reaching movements towards the opposite direction: when the perturbation was first applied, the participants under-reached the target in movements toward the non-delayed side. This initial generalization was consistent between the left and right workspace specific delay groups. However, after adaptation, no aftereffects were observed in movements toward the non-delayed side in both groups. This does not contradict our findings: in the Farshchiansadegh, participants adapted to the delay only in one workspace, and after adaptation, they were examined for aftereffects in the other workspace. In contrast, in our study, the participants adapted and examined for aftereffects in the entire workspace, but with the presence of delay in movements toward only one workspace.

We found that the effect of adaptation to a workspace-and direction-specific delay during a reaching task transferred to the blind circle drawing task. These circle-drawing movements can be considered as rhythmic movement, which are considered significantly distinct from discrete reaching movement in various aspects (Spencer et al., 2003, Buchanan et al., 2006, Hogan and Sternad, 2007). Therefore, our results are consistent with a study that showed transfer of adaptation to visuomotor delay between reaching movements to slice rhythmic movements and vice versa (Botzer and Karniel, 2013). Furthermore, transfer of adaptation to delayed visual feedback during reaching task to rhythmic movements without visual feedback was also observed (Botzer and Karniel, 2013). Our results are also in agreement with previous results that showed transfer of adaptation to visuomotor rotation during discrete reaching movements to rhythmic slice movements (Scheidt and Ghez, 2007).

It is still a matter of debate how delay is represented in the motor system. Adaptation to delayed information can be obtained by representing the perturbation as time-based or state-based. On one hand, recent studies provided support for time-based representation of both delayed force field and visual feedback (Witney et al., 1999, Levy et al., 2010, Rohde et al., 2014, Leib et al., 2015, Avraham et al., 2017). In contrast, other studies provided evidences for state-based representation, and that participants were not able to correctly represent the delay as time difference. For example, adding a delay to force feedback affects stiffness perception (Pressman et al., 2007, Nisky et al., 2011, Di Luca et al., 2011). This suggest that humans are not able to perceive the delay as time difference between the sensory inputs, and therefore, are unable to realign the different sensory inputs to avoid perceptual biases. Our results are inconsistent with time-based representation – the participants modified their movements’ extent following exposure to delay, and exhibited aftereffects when the delay was unexpectedly removed – if they would represent the time difference they would have modified the timing of movements rather than their amplitude.

Once agreed on a state-based representation, which one is used? It was suggested that the misalignment between the hand and the cursor is interpreted as a mechanical load of mass (the cursor) with a spring and a damper that connects between the hand and the cursor. This model was used to explain the changes in grip forces accompanied with delayed visual feedback (Sarlegna et al., 2010), the changes in resistive sensation following adaptation to visuomotor delay (Takamuku and Gomi, 2015), and the generalization between adapting to a visuomotor delay or to a mechanical system between the hand and the cursor (Leib et al., 2017). In addition, representation of mechanical system also explains the changes in movements after adapting to force field (Wang et al., 2001). Another possible state-based representation of visuomotor delay is considering an increase in gain between the hand and the cursor [ref-Guy]. Both mechanical system and gain representation can be used to explain the hypermetria in our results. Therefore, for simplicity of implementation and interpretation, in our computational model we used a gain representation of the delayed visual feedback. Using gain representation, we were able to simulate the results observed in our experiment both in reaching and blind drawings.

### On the other hand

It could be potentially interesting to repeat our experiments with the left hand of either right- or left-handed individuals. However, right-handed individuals use additional cognitive structures outside of the motor system to learn a motor task with the left hand (Grafton et al., 2002). Therefore, examining adaptation to delay with the left hand is not likely to provide a substantial contribution to the validation of our model. Furthermore, testing our model with left-handed participants may also be of limited value for testing our current hypotheses as there are many differences between left and right handed, as demonstrated in the evidence that the cerebral organizations of the hemispheres are not mirror images of each other (Wolff et al., 1977). Such differences were observed in the functional connectivity between motor areas in the two hemispheres in a resting state, which was significantly higher for right handed participants (Pool et al., 2015). This functional connectivity between the hemispheres in right handed may play an important role in learning lateralized perturbation such as the one presented in our study, and therefore, studying left-handed individuals while of interest, is also unlikely to validate our findings.

### Hemi-spatial neglect as a spatio-temporal deficit

Hemispatial neglect occurs more frequently and more severely after right hemisphere lesion. However, the basis for this phenomenon and for other impairments involved with unilateral brain damage is still unknown. (Heilman and Valenstein, 2003) proposed a model to explain the imbalance between the hemispheres, in light of motor impairments such as neglect. They argued that the asymmetry in attention and intention between the hemispheres is a result of asymmetrical representation of the workspaces, such that the right hemisphere incorporates representations for both workspaces, yet the left hemisphere hold representation only for the right workspace. However, in addition to the spatial deficit observed in neglect, several studies also reported time-related impairments. For example, reports of a considerable delay in visual awareness of left stimuli compared to right stimuli (Robertson et al., 1998). In addition, neglect patients present prolonged contralesional and shorter ipsilesional attentional blink – the time interval between two sequentially presented targets in which a second target can be detected (Adair and Barrett, 2008). Finally, neglect patients, compared to healthy individuals, significantly underestimate the durations of stimuli (Danckert et al., 2007). This suggests that neglect is a spatial-temporal rather than a purely spatial deficit, and that there is a link between laterality and temporal aspects of information processing. Hence, we suggest here that the imbalance between the hemispheres can also be associated with visuo-temporal processes.

Previously, by analyzing drawings performed by spatial neglect patients, studies showed a defective ability of the patients to draw the side of the object in the side contralateral to their brain lesion, resulting in asymmetrical drawing (Gainotti et al., 1972). Such asymmetry was also observed as a result of exposing participants to spatial shift by using prism adaptation, which was also proved to assist in shifting the perceived midline of neglect patients (Smith and Bowen, 1980, Adair and Barrett, 2008, Champod et al., 2016). It was suggested that the adaptation process is the fundamental cause for improving neglect symptoms. Mainly, this adaptation process stimulated areas involved in functions related to multisensory integration and spatial representation (Rossetti et al., 1998). Although prism adaptation has been shown to be effective in improving neglect symptoms, not all patients were able to adapt and benefit from this method (Serino et al., 2007). Similarly to prism adaptation, our results show that in the case of adaptation to lateralized time-based perturbation, movements become asymmetrically hypermetric. This effect on healthy participants was present even after the perturbation was removed and the aftereffects of adaptation washed out. Since the hypermetria in participants’ drawings was demonstrated in asymmetrical elongation, we believe that adaptation to visuomotor delay may be investigated as a potential method to alleviate spatial neglect symptoms with potentially long-lasting effects. However, additional studies on healthy individuals are needed to determine how long would it take for the hypermetric circles to wash out.

The observed connection between time and space, demonstrated through our model, can help explain the motor deficits observed in neglect, which has been suggested to be associated with distortions in time processing. By integrating the model for unilateral neglect with our proposed model, we can further establish the connection between temporal perturbations and spatial-motor impairments. Understanding the role of each hemisphere in mediating time and space representation can provide important insights on pathological cases involving injury in only one side of the brain and also to provide new directions for diagnosis and rehabilitation.

## Methods

### Participants and experimental setup

Eighty right-handed healthy volunteers (ages 18-35) participated in the study after signing the informed consent form as required by the Human Participants Research Committee of Ben-Gurion University of the Negev, Be’er-Sheva, Israel. The participants were all naive to the purpose of the experiment and were paid to participate.

The experiment was administered in a virtual reality environment in which the participant held a robotic arm: six degrees of freedom (DOF) PHANTOM^®^ Premium^TM^ 1.5 haptic device (Geomagic^®^), controlled by a dedicated C++ code. Participants held the robotic arm with their right hand controlling a cursor displayed on a screen and aligned with their hand location. Participants’ hand was hidden from sight the entire experiment by the screen that was located horizontally above their hand, and by a sheet that covered their upper body. Hand movements were constrained to the horizontal plane by an air sled wrist-supporter that reduces friction with the surface. The update rate of the control loop was 1000Hz.

### Protocol

The experiment consisted of two tasks: reaching movements to left or right targets and circle drawing without visual feedback. The trials were presented in a random and predetermined order between participants. In the reaching task, a trial was initiated when participants placed a circular cursor, 1 cm diameter, inside a starting point with the same size. The task was to move the cursor from the starting point to a circular target, 2 cm diameter, which appeared in the left or the right side of the task space, at a distance of 10 cm away from the starting position (Figure 2). Movement started when the color of the cursor changed, instructing the participant to perform a smooth point-to-point reaching movement. Movement ended when the velocity was less than 1 cm/s. Following the movement, the robot applied a spring-like force that returned the hand to the start position. In addition, the participant received a feedback based on the velocity of the movement and on the accuracy. When the maximum velocity was lower than 30 cm/s, the word “Faster” appeared on the screen, and when the velocity was higher than 50 cm/s, the word “Slower” was displayed. Accurate movements were defined as those in which the center of the cursor was in the range of ±1 cm from the center of the target. To provide a feedback about the end movement position, we presented the location of the cursor with a color cue that indicated the accuracy of the movement (green for accurate movement and red for inaccurate movement).

In the circle drawing task, a circle with radius of 3.5 cm was displayed on the screen in four different locations: front, back, right and left. Arrows on the circle indicated the direction of the drawing to either clockwise or counterclockwise. The location of the starting point was always in the middle of the task space in all conditions, identically to the location of the start point in the reaching task. A trial was initiated when participants placed a circular cursor, 1 cm diameter, inside the starting point. When the cursor disappeared and the start point changed its color, the participants were required to initiate a smooth circular movement along the desired circle from the starting point, in the direction of the arrows. Movement ended when the velocity was less than 0.5 cm/s.

Participants were assigned to one of four groups according to the workspace where they were exposed to delay: (1) only in leftward reaching movements (Left Delay, N=15), (2) only in rightward reaching movements (Right Delay, N=15), (3) in both leftward and rightward movements (Both Delay, N=20), and (4) a control group that was not exposed to any perturbation throughout the entire experiment (No Delay, N=15). The first block of the experiment (40 trials) was training for the circle drawing task. In these training trials, participants drew the circles without visual feedback. After each trial, the drawn circle was displayed along with the desired circle and the start point. The purpose of these trials was to acquaint the participants with the task and to train them to draw circles according to a desired trajectory when no visual feedback is presented. The data from the training trials were not included in data analysis. Then, the experiment was divided into three sessions: Baseline, Adaptation and Washout. In the Baseline session (160 reaching movements and 40 circle movements), participants performed reaching without any perturbation and with interleaved blind circle-drawings. After the baseline session, we presented participants with another block of training for the circle drawing task (16 trials). The purpose of this block was to verify that the circles drawn in the Adaptation session originated from the exposure to the applied perturbation and not from forgetting how to draw the blind circles. In the adaptation session (416 reaching movements and 104 circle movements), the visual feedback between the hand and the cursor in the reaching task was delayed by 0.15 sec either when the left target appeared (LD), when the right target appeared (RD), or when both right and left targets appeared (BD), depending on the experimental group. For the No-Delay group, there was no change in the Adaptation session. During Washout (160 reaching movements and 40 circle movements), the delay was unexpectedly removed, which enabled us to examine the aftereffect of adaptation. The entire experiment lasted approximately 90 minutes with four breaks of 1.5 minutes every 160 reaching trials.

### Data analysis

Position and velocity were recorded during the entire experiment at 200Hz and were analyzed off-line using custom-written Matlab® code (The MathWorks, Inc., Natick, MA, USA). Both position and velocity were filtered by low pass Butterworth filter with a cutoff frequency of 10Hz (Matlab function `filtfilt()`). In addition, the position was interpolated to fit the number of samples using Matlab function `interpft()`, which resulted in different sampling rate for each signal that depended on the number of samples in the original signal. For the purpose of data analysis, we defined reach movement initiation when the velocity rose above 5% of the maximum velocity, and movement ending when the velocity decreased below 5% of the maximum velocity. We examined the trajectory in each direction separately, by measuring the amplitude of the movement as the maximum displacement.

In the circle drawing task, due to the importance of the drawing’s direction in our study, we first removed all circles that were mistakenly drawn in the direction that was opposite to the instructed direction (1.65% of all circles). Then, we defined the initiation and end of the movement by using both position and velocity. Initially, we found the locations where the hand first leaves and returns to the start position area. This was done to account for only one circle in cases when the participants drew more than one complete circle. Afterwards, we defined the actual initiation and end of the movement based on the velocity thresholds that we defined in the reaching movements. To calculate the deviation of the drawn circles from the desired circle, we measured the peak amplitude of hand movement in the x and y directions.

In the analysis of the drawn circles, we did not include the data from the Early Adaptation stage. From the results of the reaching task in all the conditions, we saw that participants adapted to the perturbation quite fast. Therefore, we could not verify that all drawn circles in all 8 conditions, used for the analysis, are in this phase of post-exposure and pre-adaptation.

### Simulation of arm movement

Aiming to examine whether our results can be explained by the mechanics of arm movement solely, we modeled arm dynamics as two link model with two variables of shoulder 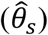 and elbow 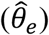 configuration. We simulated a simplified control of arm movement by two controllers of trajectory and end-point. This model was used to simulate both reaching movements and blind circular movements. To simulate reaching movements, we assumed that the controller tracks a planned minimum-jerk trajectory defined as a smooth trajectory from start to end-position along the x-axis (Flash and Hogan, 1985). Desired circular movements were defined by fitting a 12^th^ order polynomial function to a desired trajectory, in order to achieve smooth velocity and acceleration along with the desired path. The torques required to perform a desired movement were computed from equation (1).

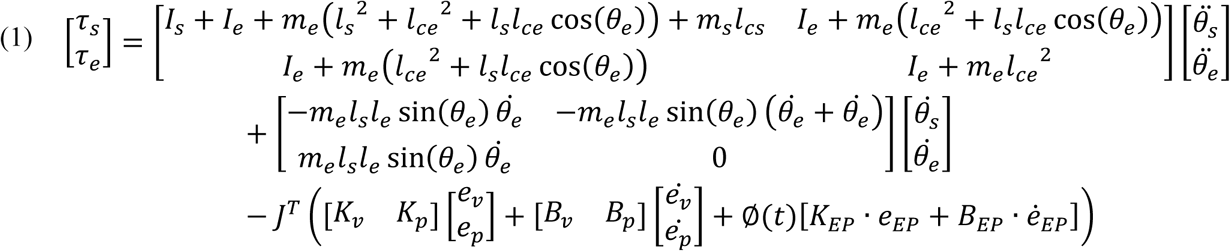

Dynamics equation of arm movement-torques are computed using two links robotics dynamics with additional PD controllers for proprioceptive and visual feedback, and for end-point controller. Values of arm parameters are similar to those used in (Scheidt and Ghez, 2007). The position (velocity) error is defined as the difference between the actual to the desired arm position (velocity). The values of all proportional (K) and derivative (B) controllers are presented in table S1 in supplemental information.

The trajectory controller consisted of two mechanisms of feedback and feedforward, utilized to execute a smooth movement along a desired path (Botzer and Karniel, 2013). We assumed that the feedforward controller is an inverse model of the forward model. Therefore, any changes in the inverse model as a result of adapting to a new dynamics in the environment will also lead to changes in the relevant forward controller. By using two different feedforward and feedback controllers for vision and proprioception, we were able to differentiate between movements with and without visual feedback.

The end-point controller did not change throughout the simulation, and was always used to stabilize the hand at the desired end-position. This controller was multiplied by a sigmoid function Ø*(t)*, which increased the contribution of the end-point controller according to a desired timing along the movement. Before adapting to the delay in both the reach and circular movements, the time when the sigmoid function was equal to 0.5 was at the end of movement. After adapting to the delay, we assumed that as a result of uncertainty during the movement, the end-point controller would be tuned earlier – approximately in the middle of the movement.

The different stages in the experiment were simulated by changing the visual delay and the magnifying gain that represented the adaptation to delay. This way we could distinguish between the different stages and simulate the different movements observed in the experiment. In the Late Baseline, the participants were not exposed to the delay yet (ΔT_v_=0), and no adaptation process has occurred (G=1). In the Early Adaptation, the visual delay was set to ΔT_v_=0.15 sec, and the gain still did not change (G=1). After adapting to the delay (Late Adaptation), the gain was changed to G=1.2 such that the desired trajectory was extended in the direction of the movement. In this stage, the visual delay was ΔT_v_=0.15 sec. To simulate the removal of the delay in the Early Washout stage, the visual delay was changed to ΔT_v_=0, and the gain in this stage was G=1.2. Throughout the experiment, the proprioception delay was not change, and therefore we set ΔT_p_=0.

### Statistical analysis

The effect of the perturbation in each condition on the reaching movements was assessed by using a two-way repeated measures ANOVA with between factors of Stage (LB/EA/LA/EW) and Direction (Leftward Movements/Rightward Movements). For the blind drawings, we initially examined the effect of delay on left and right error separately, using one-way repeated measures ANOVA with factor Stage (LB/LA/EW). After dividing between the circles according to initiation workspace, the lateral effect on the blind drawings was examined using two-way repeated measures ANOVA with within factors of Stage (LB/LA/EW) and Initiation-workspace (Left/Right). Data were tested for normality distribution using Lilliefors test. Additionally, we used Mauchly’s test to examine whether we can assume sphericity of the data. In case the sphericity assumption was not met, we used Greenhouse-Geisser adjustment. When found significant effects, post-hoc t-test was performed with the Bonferroni correction. Significant effects were defined at the p<0.05 probability level.

## Supplemental Information

### Circle drawing task

We calculated the deviation of the circles in y-axis (toward front and back directions). We found no similar pattern of deviation as in x-axis (Figure S1 related to figures 4C-F and 6), which indicate that the imposed perturbation caused only deviation toward the left and right workspaces. However, from examining our experimental setup, we found that movements toward front and back directions were partly constrained because of space limitations. Therefore, in order to fully assess the effect of workspace-specific delay on movements in these directions, further experiments are required.

**Figure S1.**
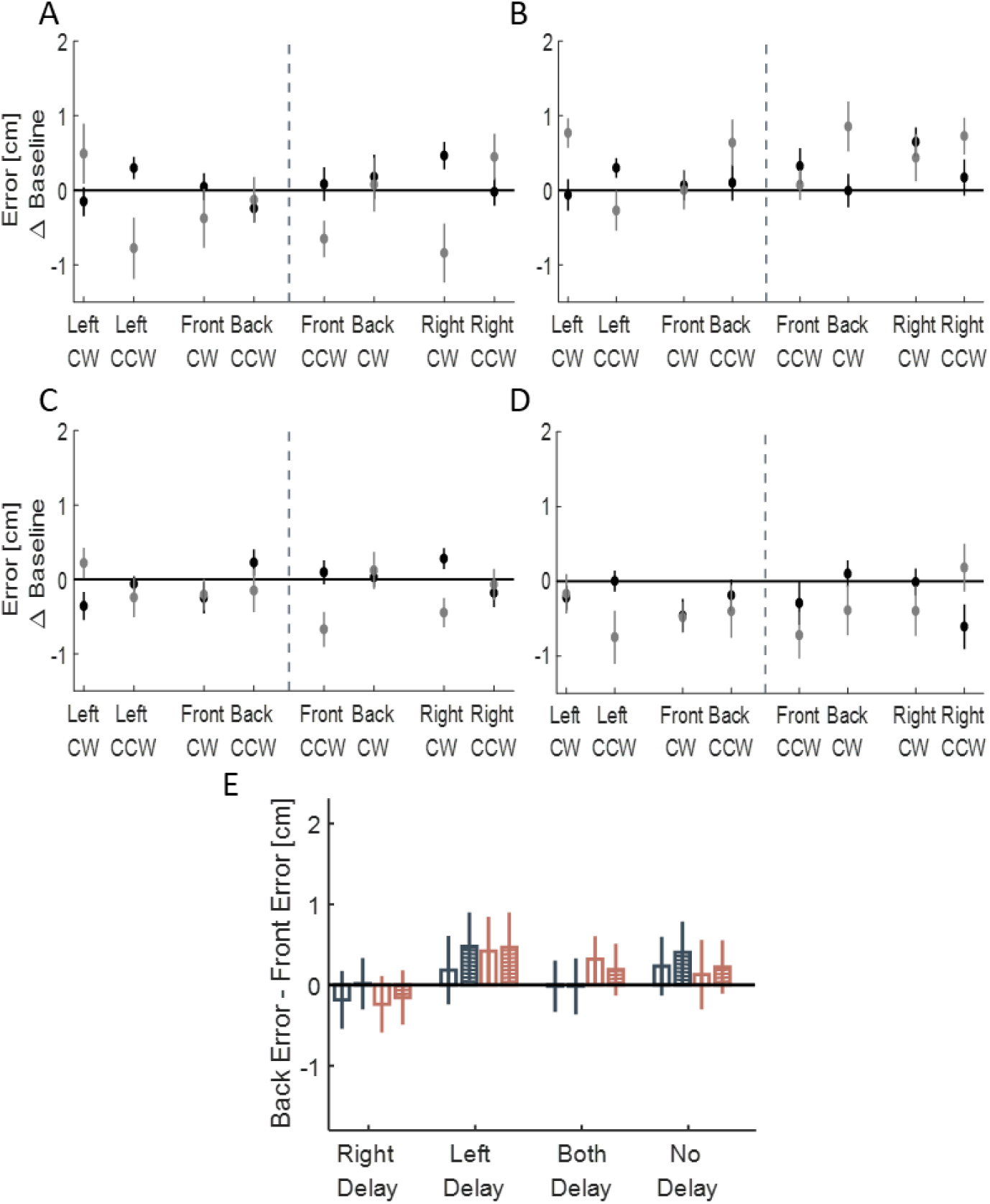
Deviation of the drawn circles toward front (gray) and back (black) directions, divided to left and right initiation (dashed line), for (A) Left delay group, (B) Right delay group, (C) Both delay group and (D) No delay group. The X-axis represents the location of the drawn circle (front, back, right and left) and the direction of clockwise (CW) and counterclockwise (CCW). No pattern of deviation is observed in those directions. (E) Statistical analysis of Back error - Front error Difference for the circles initiated in the left (dark blue) and in the right (light red). Empty bars are for Late Adaptation session and bars with stripes are for Washout session. From the graph, no similar pattern of elongation toward front or back is observed.

### Simulation study

To simulate arm dynamics, we used mathematical model of planar two-link kinematic chain. The dynamic equation of the system was written as

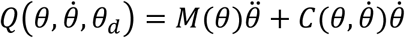

where 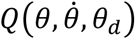 is the joints torques generated by the controller, *θ* is a vector of shoulder and elbow joints angles, *M*(*θ*) is the inertia matrix, and 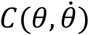 is the Coriolis and centripetal coefficients matrix. The controllers’ equation predicted the outcome of a motor command by using inverse model of arm dynamics (feedforward controller) and proportional-derivative controller (feedback controller). Accordingly the controller was calculated using

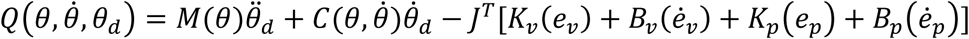

where *K* and *B* are proportional and derivative coefficients of the PD controller, separated for vision and proprioception. *e* and 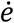 represents the error between the actual to the desired arm position and velocity respectively. The feedback was calculated in Cartesian coordinates, and therefore, was multiplied by the transpose Jacobean (*J*^*T*^) to transform to joints coordinates.

Additionally, we added end-point controller for stabilization on the desired end point. The values of the controllers used in our simulation are depicted in table S1 (related to equation 1 and figure 7). Using the computational model, we were only partially able to simulate the results observed in our experiment. Therefore, to accurately simulate the asymmetrical results, a further *Attentional-Motor Asymmetry* model for the effect of delay on the hemispheres was required (see results section for more details).

**Table S1.** Values of proportional and derivative controllers

| Parameter | Value |
| --- | --- |
| $K_p$ | 0.9 N/m |
| $B_p$ | 0.7 N·s/m |
| $K_v$ | 0.9 N/m |
| $B_v$ | 0.7 N·s/m |
| $K_{EP}$ | 0.9 N/m |
| $B_{EP}$ | 0.7 N·s/m |
| Visual Gain ($G_v$) | 0.7 |
| Perception Gain ($G_p$) | 0.3 |
| Adaptation Gain ($G$) | 1.2 |

